# Absorption changes in Photosystem II in the Soret band region upon the formation of the chlorophyll cation radical [P_D1_P_D2_]^+^

**DOI:** 10.1101/2022.05.12.491653

**Authors:** Alain Boussac, Miwa Sugiura, Makoto Nakamura, Ryo Nagao, Takumi Noguchi, Stefania Viola, A. William Rutherford, Julien Sellés

## Abstract

Flash-induced absorption changes in the Soret region arising from the [P_D1_P_D2_]^+^ state, the chlorophyll cation radical formed upon light excitation of Photosystem II (PSII), were measured in Mn-depleted PSII cores at pH 8.6. Under these conditions, Tyr_D_ is *i*) reduced before the first flash, and *ii*) oxidized before subsequent flashes. In wild-type PSII, when Tyr_D_^●^ is present, an additional signal in the [P_D1_P_D2_]^+^-*minus*-[P_D1_P_D2_] difference spectrum was observed when compared to the first flash when Tyr_D_ is not oxidized. The additional feature was “W-shaped” with troughs at 434 nm and 446 nm. This feature was absent when Tyr_D_ was reduced, but was present *i*) when Tyr_D_ was physically absent (and replaced by phenylalanine) or *ii*) when its H-bonding histidine (D2-His189) was physically absent (replaced by a Leucine). Thus, the simple difference spectrum without the double trough feature at 434 nm and 446 nm, seemed to require the native structural environment around the reduced Tyr_D_ and its H bonding partners to be present. We found no evidence of involvement of P_D1_, Chl_D1_, Phe_D1_, Phe_D2_, Tyr_Z_, and the Cyt*b*_559_ heme in the W-shaped difference spectrum. However, the use of a mutant of the P_D2_ axial His ligand, the D2-His197Ala, shows that the P_D2_ environment seems involved in the formation of “W-shaped” signal.

## Introduction

Oxygenic photosynthesis is responsible for most of the energy input to life on Earth. This process converts the solar energy into fiber, food and fuel, and occurs in cyanobacteria, algae and plants. Photosystem II (PSII), the water-splitting enzyme, is at the heart of this process, see (Cox et al. 2020) for a recent review.

Mature cyanobacterial PSII generally consists of 20 subunits with 17 trans-membrane and 3 extrinsic membrane proteins. PSII binds 35 chlorophylls *a* (Chl-*a*), 2 pheophytins (Phe), 1 membrane b-type cytochrome, 1 extrinsic c-type cytochrome, 1 non-heme iron, 2 plastoquinones (Q_A_ and Q_B_), the Mn_4_CaO_5_ cluster, 2 Cl^−^, 12 carotenoids and 25 lipids (Suga et al. 2015). The 4^th^ extrinsic PsbQ subunit was also found in PSII from *Synechocystis* sp. PCC 6803 in addition to PsbV, PsbO and PsbU (Gisriel et al. 2022).

Among the 35 Chl’s, 31 are antenna Chl’s and 4 (P_D1_, P_D2_, Chl_D1_ and Chl_D2_), together with the 2 Phe molecules, constitute the reaction center pigments of PSII. After the absorption of a photon by the antenna, the excitation energy is transferred and trapped in the reaction center (Mirkovic et al. 2017). After a few picoseconds, a charge separation occurs resulting in the formation of the Chl_D1_^+^Phe_D1_^−^ and then of the [P_D1_P_D2_]^+^Phe_D1_^−^ radical pair states, with the positive charge mainly located on P_D1_, *e.g.* (Capone et al. 2023, Holzwarth et al. 2006, Romero et al. 2017). The main PSII cofactors are shown in Fig. 1 for PSII with PsbA3 as the D1 protein (Nakajima et al. 2022).

**Figure 1:**
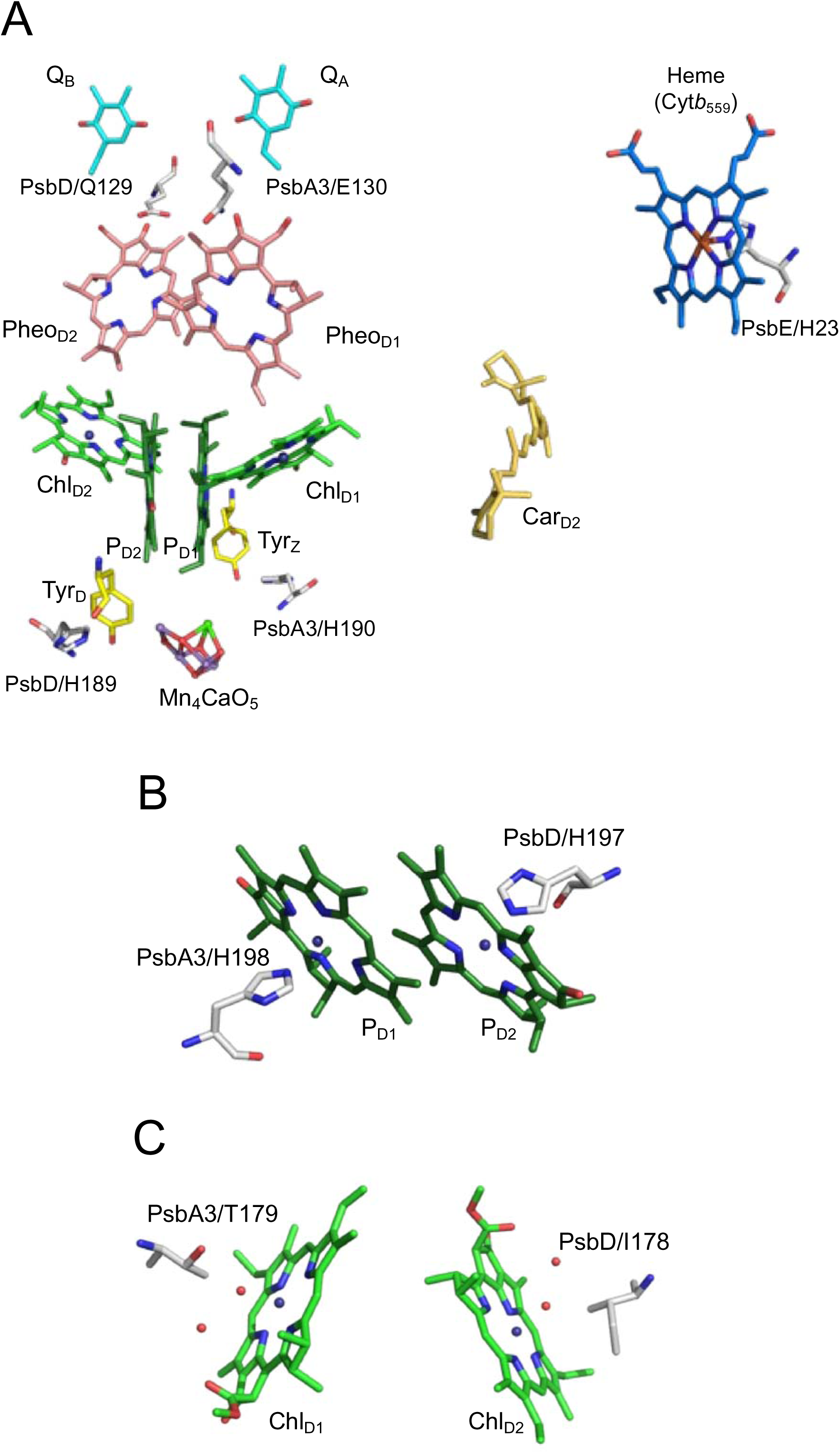
Panel A: Arrangement of cofactors of the *T. elongatus* PSII involved in the electron transfers and studied in the present work together with some amino acids interacting with these cofactors. Panel B, structure of P_D1_ and P_D2_ with their amino acid residue ligands. Panel C, structure around Chl_D1_ and Chl_D2_. The PsbA3/E130 residue is H-bonded to Phe_D1_ and PsbD1/Q129 residue is H-bonded to Phe_D2_. The figures were drawn with MacPyMOL with the A monomer in PDB 7YQ7 (Nakajima et al. 2023).

After the formation of [P_D1_P_D2_]^+^Phe_D1_^−^, the electron on Phe_D1_^−^ is transferred to Q_A_, the primary quinone electron acceptor, and then to Q_B_, the second quinone electron acceptor. While Q_A_ is only singly reduced under normal conditions, Q_B_ accepts two electrons and two protons before leaving its binding site and being replaced by an oxidized plastoquinone molecule from the membrane plastoquinone pool (Boussac et al. 2010, de Causmaecker et al. 2019, Fufezan et al. 2005, Sedoud et al. 2011 and references therein). On the donor side of PSII, [P_D1_P_D2_]^+^ oxidizes Tyr_Z_, the Tyr161 of the D1 polypeptide. The Tyr_Z_^●^ radical is then reduced by the Mn_4_CaO_5_ cluster, *e.g.* (Lubitz et al. 2019, Shevela et al. 2023) for some reviews. After four charge separations, the Mn_4_CaO_5_ cluster accumulates four oxidizing equivalents and thus cycles through five redox states denoted S_0_ to S_4_. Upon formation of the S_4_-state, two molecules of water are oxidized, the S_0_-state is regenerated and O_2_ is released (Joliot et al. 1969, Kok et al. 1970).

In O_2_ evolving PSII with an intact Mn_4_CaO_5_ cluster, the electron transfer from Tyr_Z_ to [P_D1_P_D2_]^+^ takes place in time ranges from tens of ns to tens of µs, *e.g.* (Renger 2012). The kinetics in the ns range are sometimes discussed in term of a pure electron transfer, whereas in the µs range they involve large proton relaxations in the H-bond network, *e.g.* (Renger 2012). The fast electron transfer from Tyr_Z_ to [P_D1_P_D2_]^+^ occurs with a *t*_1/2_ close to 20 ns (Brettel et al. 1984, Gerken et al. 1987). The µs to tens of µs phases correspond to proton movements in which the phenolic proton of the Tyr_Z_^●^ radical moves, in a first step, onto the H-bonded His190 of the D1 polypeptide.

In Mn-depleted PSII, the electron transfer from Tyr_Z_ to [P_D1_P_D2_]^+^ is strongly pH dependent (Conjeaud and Mathis 1980) with a *t*_1/2_ of ~ 190 ns at pH 8.5 and ~ 2-10 µs at pH 6.5 (Faller et al. 2001). At pH 8.5 and above, Tyr_D_, the Tyr160 of the D2 polypeptide that is symmetrically positioned with respect to Tyr_Z_ (Fig. 1), is slowly reduced in the dark (Boussac and Etienne 1982) and becomes able to donate an electron to P_D1_^+^ with a *t*_1/2_ of ~ 190 ns, similar to Tyr_Z_ donation (Faller et al. 2001). Consequently, at room temperature and pH 8.5, the reduced Tyr_D_ can donate an electron to [P_D1_P_D2_]^+^ in approximately half of the centers upon a saturating ns flash in Mn-depleted PSII, while in the remaining half of the centers [P_D1_P_D2_]^+^ is reduced by Tyr_Z_. Based on the electron transfer rate and the distance between TyrD and PD1 and PD2, it is considered that Tyr_D_ oxidation occurs by donation to P_D2_^+^, which shares the cation with P_D1_ by a redox equilibrium, [P_D1_^+^P_D2_] ↔ [P_D1_P_D2_^+^], (Rutherford et al. 2004).

The difference spectra [P_D1_P_D2_]^+^-*minus*-[P_D1_P_D2_] measured in the Soret region of the Chl absorption, and corrected for the contribution of Q_A_^−^-*minus*-Q_A_ formation, shows a large bleaching at ~ 432 nm in wild type PSII (Diner et al. 2001). Additionally, the difference spectrum exhibits a negative feature between 440 nm and 460 nm. The origin of this additional spectral feature that is observed in both inactive PSII (Diner et al. 2001) and O_2_-evolving PSII, *e.g.* (Sugiura et al. 2004), has not been determined. In this spectral region, the redox changes of several species, such as the Chls, cytochromes and amino acid radicals may give rise to absorption changes upon a charge separation event. However, the difference spectra recorded at time as short as 20 ns after an actinic ns laser flash and decaying with the same kinetics as [P_D1_P_D2_]^+^ cannot originate from states other than those with [P_D1_P_D2_]^+^ present. Alternatively, band shifts of either Chl’s or cytochromes or amino acid radical(s) induced by the formation of [P_D1_P_D2_]^+^ may also contribute.

In the present work, we have recorded the [P_D1_P_D2_]^+^Q_A_^−^-*minus*-[P_D1_P_D2_]Q_A_ difference spectra in PSII from different cyanobacteria species and mutants (see Fig.1) in order to clarify the nature of the spectral feature between 440 nm and 460 nm. All the measurements were done in Mn-depleted PSII after a long dark-adaptation in order to measure the [P_D1_P_D2_]^+^Q_A_ ^−^-*minus*-[P_D1_P_D2_]Q_A_ difference spectra with Tyr_D_ either reduced or oxidized, *i.e.* in the presence of Tyr_D_^●^.

## Materials and Methods

### PSII samples

The PSII samples used in this study were purified from *i*) *Thermosynechoccocus elongatus* (see below), *ii*) *Chroococcidiopsis thermalis* PCC7203 grown under far-red light (750 nm, LED750-33AU from Roithner LaserTechnik), and from the D2/H197A mutant in *<u>Synechocystis</u>* <u>PCC 6803 (Hayase et al. 2023)</u>. PSII from *C. thermalis* has 4 Chl-*f* and 1 Chl-*d* replacing 5 of the 35 Chl-*a* (Nürnberg et al. 2018, Judd et al. 2020) see also (Chen et al. 2010, Gan et 2014, Gisriel et al. 2022b).

The *T. elongatus* strains used in this study, listed in Table 1, were constructed from a strain with a His_6_-tag on the carboxy terminus of CP43, called 43H. The mutants in the D2 protein were constructed in the *psb_D1_* gene with the *psb_D2_* gene deleted.

**Table 1:**
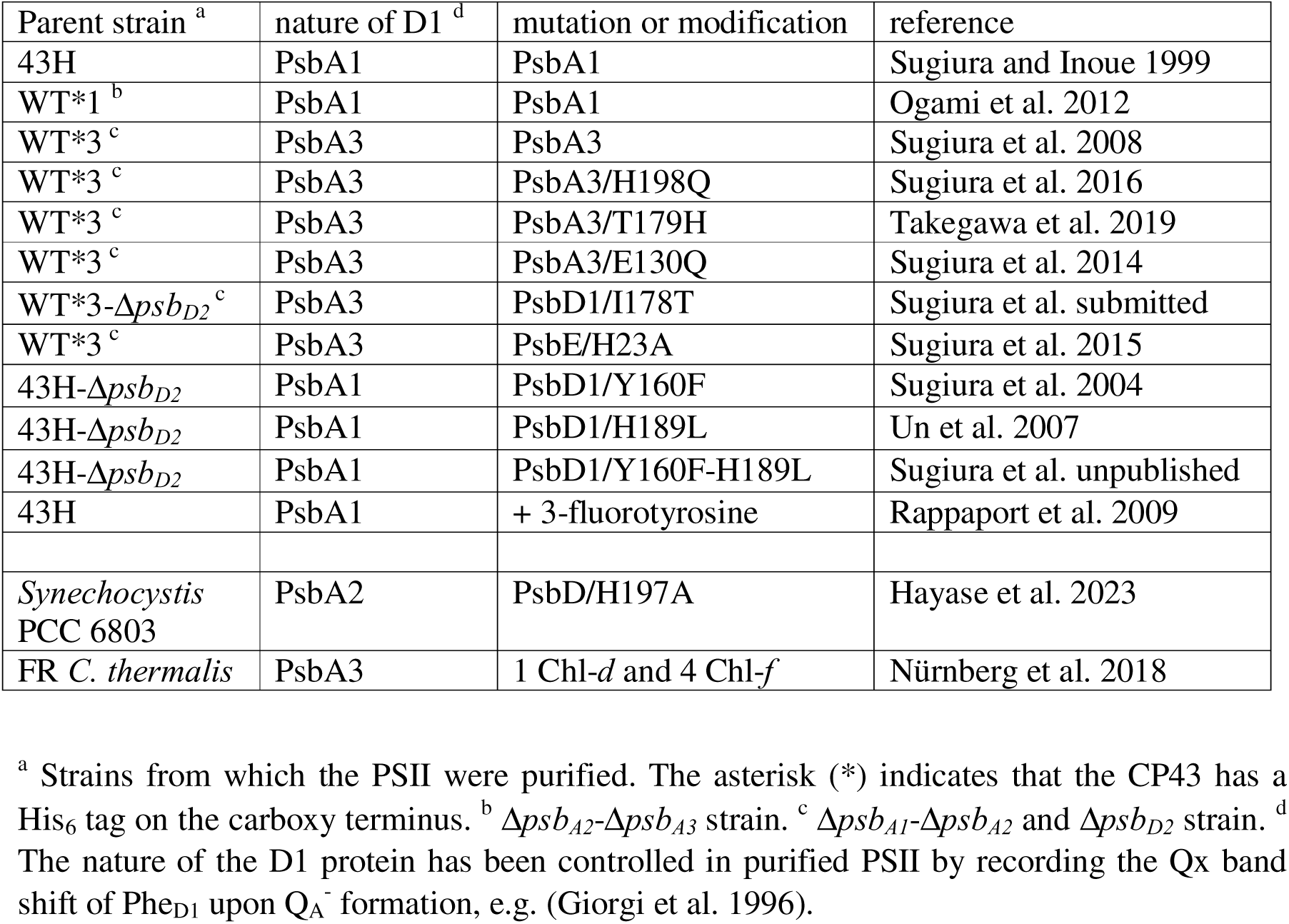
PSII samples used in this study.

PSII purifications from the *T elongatus* strains were performed as previously described (Sugiura et al. 2014). For Mn-depletion, 20 mM NH_2_OH, from a stock at 1 M at pH 6.5, and 1 mM EDTA were added to the PSII samples. After an incubation for approximately 1 minute in dim light in the cold room at 4°C, the hydroxylamine and EDTA were removed by washings of the PSII samples by cycles of dilution (taking each approximately 30 minutes) in 1 M betaine, 15 mM CaCl_2_, 15 mM MgCl_2_, 40 mM MES, pH 6.5, followed by concentration using Amicon Ultra-15 centrifugal filter units (cut-off 100 kDa) until the estimated residual NH_2_OH concentration was lower than 0.1 μM in the concentrated PSII samples before the final dilution for the ΔI/I measurements. In the final dilution step, the PSII samples were suspended in 1 M betaine, 15 mM CaCl_2_, 15 mM MgCl_2_, 100 mM Tris, pH 8.6.

The PSII from *C. thermalis* grown under far-red light was purified as previously described (Nürnberg et al. 2018), and then also treated with NH_2_OH as described above. The D2/H197A with a His-tag was purified with the same protocol as *T. elongatus* except the breaking of the cells with the French press that was done after resuspension of the cells in 100 mM Tris pH 8.0.

### UV-visible time-resolved absorption change spectroscopy

Absorption change measurements were performed with a lab-built spectrophotometer (Béal et al. 1999) in which the absorption changes were sampled at discrete times after the actinic flash by short analytical flashes. These analytical flashes were provided by an optical parametric oscillator (Horizon OPO, Amplitude Technologies) pumped by a frequency tripled Nd:YAG laser (Surelite II, Amplitude Technologies), producing monochromatic flashes (355 nm, 2 nm full-width at half-maximum) with a duration of 5 ns. Actinic flashes, for all the samples studied, were provided by a second Nd:YAG laser (Surelite II, Amplitude Technologies) at 532 nm, which pumped an optical parametric oscillator (Surelite OPO plus) producing monochromatic saturating flashes at 695 nm with the same pulse-length. The two lasers were working at a frequency of 10 Hz and the time delay between the laser delivering the actinic flashes and the laser delivering the detector flashes was controlled by a digital delay/pulse generator (DG645, jitter of 1 ps, Stanford Research). The path-length of the cuvette was 2.5 mm.

For the ΔI/I measurements, the Mn-depleted PSII samples were diluted in a medium with 1 M betaine, 15 mM CaCl_2_, 15 mM MgCl_2_, and 100 mM Tris with the pH adjusted with HCl at pH 8.6. All the PSII samples were dark-adapted for ~ 3-4 h at room temperature (20– 22°C) before the addition of 0.1 mM phenyl *p*–benzoquinone (PPBQ) dissolved in dimethyl sulfoxide. In all cases, the chlorophyll concentration of the samples was ~ 25 µg of Chl mL^−1^. After the ΔI/I measurements, the absorption of each diluted batch of samples was precisely controlled to avoid errors due to the dilution of concentrated samples and the ΔI/I values shown in the figures were normalized to *A*_673_ = 1.75, with ε ~ 70 mM^−1^·cm^−1^ at 674 nm for dimeric *T. elongatus* PSII (Müh and Zouni 2005).

## Results and Discussion

Fig. 2 shows the observation that triggered this study. In this experiment, the ΔI/I was measured from 401 to 457 nm, 20 ns after each saturating ns laser flash of a series of 10 fired with an interval of 2 s. The material was a Mn-depleted PsbA3/PSII which was dark adapted for ~ 3-4 h at pH 8.6. This long dark-incubation allows Tyr_D_ to be reduced in the great majority of the centers (Boussac and Etienne 1982, Faller et al. 2001). The black spectrum was recorded after the first flash, *i.e.* it corresponds to the formation of the [P_D1_P_D2_]^+^Q_A_^−^ state when Tyr_D_ is not yet oxidized. The red spectrum is an average of the ΔI/I measured from the 5^th^ to 10^th^ laser flash illumination, *i.e.* it corresponds to the formation of the [P_D1_P_D2_]^+^Q_A_^−^ state in the presence of Tyr_D_^●^ after it is formed on the first actinic flashes.

**Figure 2:**
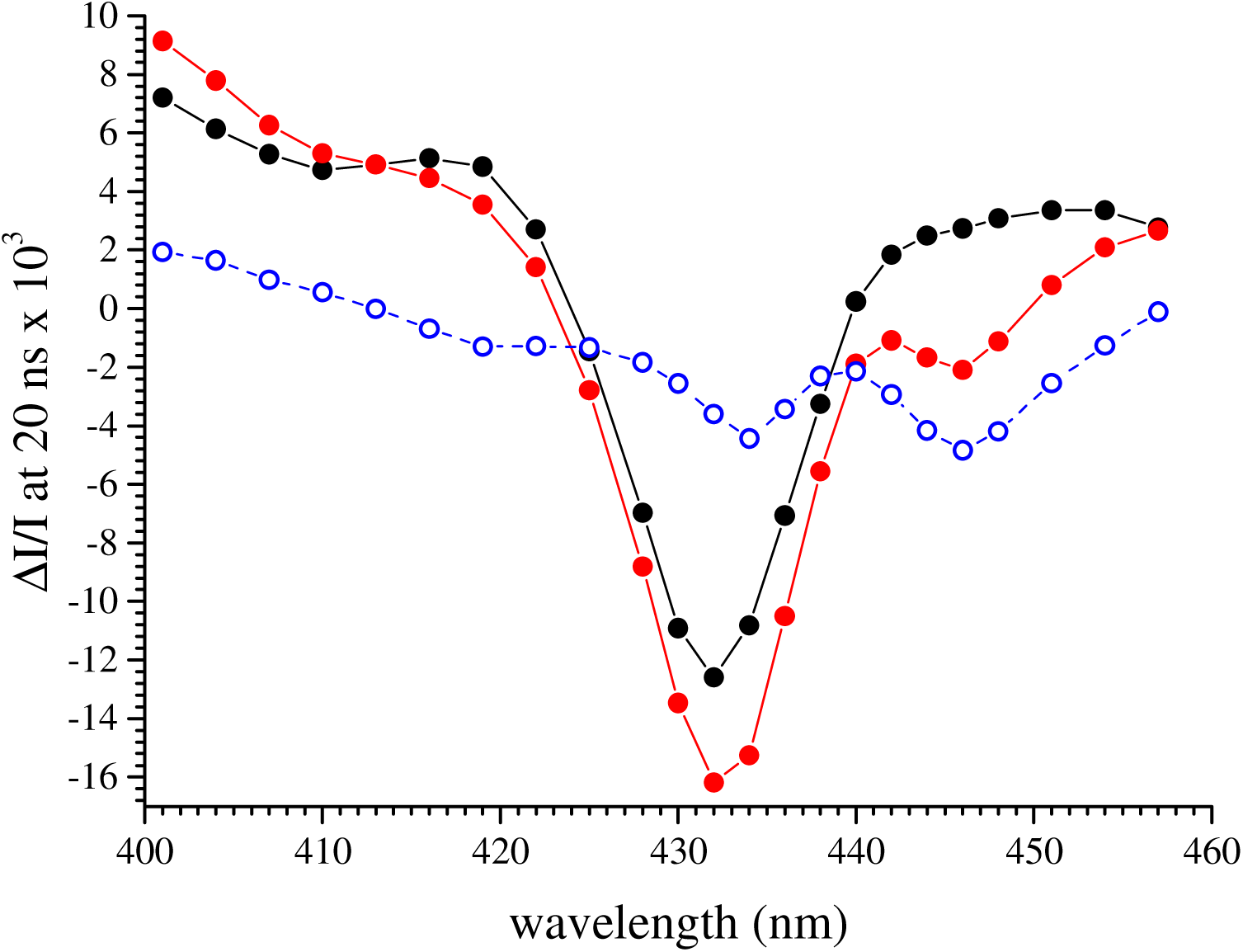
Spectra recorded 20 ns after a laser flash illumination in Mn-depleted PsbA3-PSII which was dark-adapted for 3-4 hours at pH 8.6 to allow the reduction of Tyr_D_. The black full circles were recorded after the 1^st^ flash of a sequence of 10 flashes fired 2 s apart. The red full circles are the average the measurements done from the 5^th^ to the 10^th^ flash. The blue spectrum is the red spectrum *minus* the black spectrum. The Chl concentration was 25 µg mL^−1^ and 100 µM PPBQ was added before the measurements.

In Fig. 2, the maximum bleaching in the two spectra is at 432 nm as expected for the [P_D1_P_D2_]^+^-*minus*-[P_D1_P_D2_] difference spectrum (Diner et al. 2001). However, the two spectra differ significantly. The blue spectrum, which is the red spectrum *minus* the black spectrum, shows that, once Tyr_D_^●^ has been formed after the first flashes, the [P_D1_P_D2_]^+^Q_A_^−^-*minus*-[P_D1_P_D2_]Q_A_ difference spectrum exhibits a “W-shaped” spectrum with two additional negative features, with troughs at ~ 434 nm and ~ 446 nm. This difference spectrum was calculated assuming that the same amount of [P_D1_P_D2_]^+^ was formed after the 1^st^ flash and the followings which is very likely to be the case in this sample. The very small negative contribution at 446 nm in the black spectrum is most likely due to the small fraction of centers in which Tyr_D_^●^ remained after the dark-adaptation. The trough at 434 nm in the blue spectrum could arise from the bleaching of the radical cation with a higher proportion of P_D2_^+^ in [P_D1_P_D2_]^+^. However, this would imply a difference spectrum with a more derivative shape with a contribution of the band which disappears, something that is not observed. The difference has therefore a more complex origin.

Panel A in Fig. 3, shows the results of the same experiments as in Fig. 2 but using a Mn-depleted PsbA1-PSII. The results are very similar to those in Mn-depleted PsbA3-PSII, with the additional feature in the red spectrum presenting a trough at 446 nm. The major bleaching peaks at ~ 434 nm in the red spectrum, showing that the two troughs in the blue spectrum of Fig. 2 are also present here. The similarities of these results with those in Fig. 2 shows that the nature of PsbA, *i.e.* PsbA1 *vs* PsbA3, does not affect the formation and spectrum of the additional “W-shape” structure observed after the 5^th^ to 10^th^ flashes. In Fig. 3A, a small negative contribution at 446 nm in the black spectrum (*i.e.* after the 1^st^ flash) is detected and this contribution is larger than that in Fig. 2. This difference reflects a slight variation in the proportion of centers in which the oxidized Tyr_D_ was still present upon the dark adaptation. This proportion is very weak but may vary from sample to sample.

**Figure 3:**
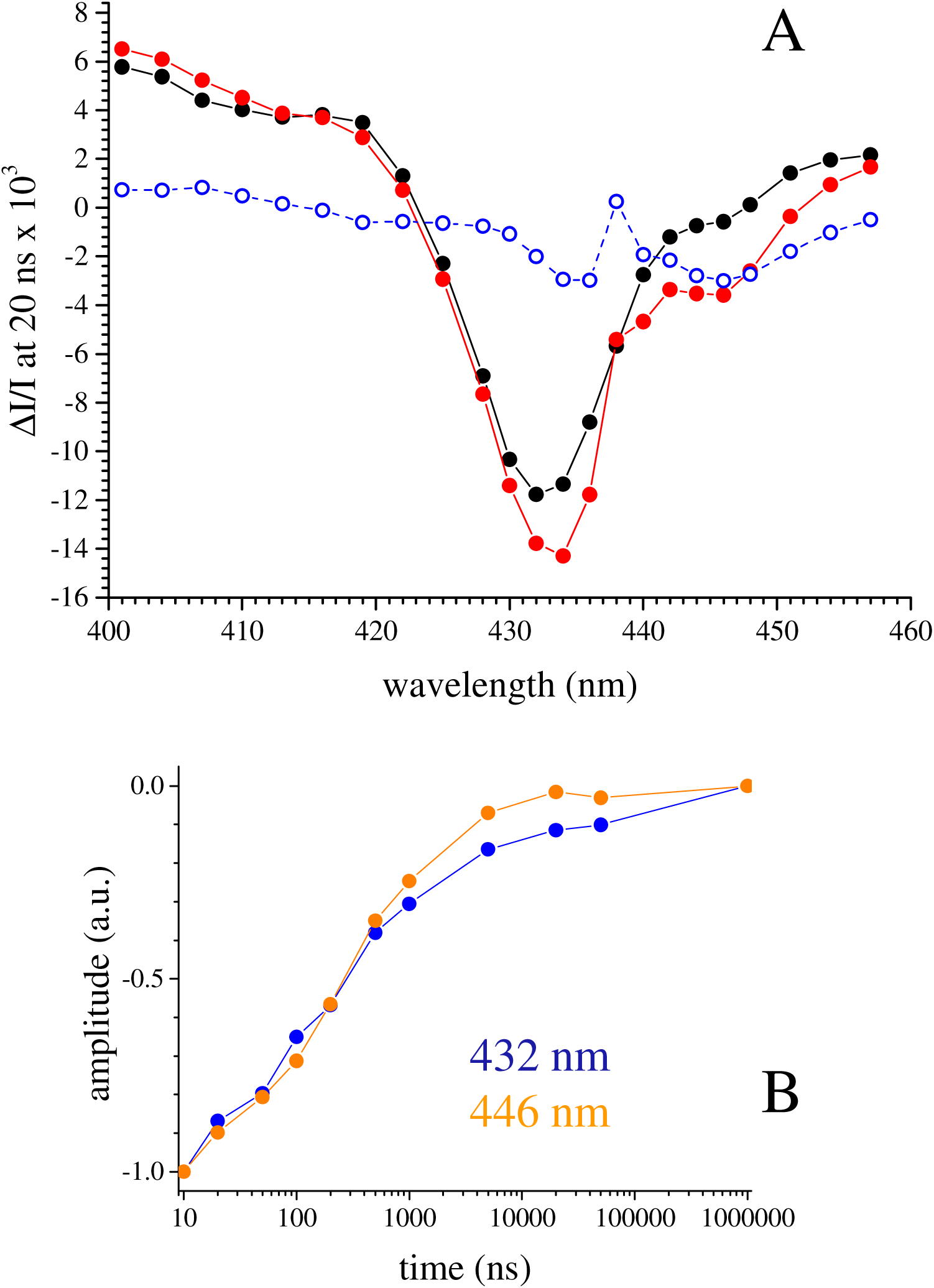
Panel A, spectra recorded 20 ns after a laser flash illumination in Mn-depleted PsbA1-PSII which was dark adapted for 3-4 hours at pH 8.6 to allow the reduction of Tyr_D_. The black full circles were recorded after the 1^st^ flash and the red full circles are the average the measurements done from the 5^th^ to the 10^th^ flash. The blue spectrum is the red spectrum *minus* the black spectrum. Panel B, decay kinetics measured from 10 ns to 1 ms in Mn-depleted PsbA1-PSII, either at 446 nm (orange) or at 432 nm (blue). The data points are averaged from the 5^th^ to the 10^th^ flash, *i.e.* when all Tyr_D_^●^ was formed in all centers. The amplitude of the signal at 10 ns was normalized to −1. The Chl concentration was 25 µg mL^−1^ and 100 µM PPBQ was added before the measurements.

Panel B in Fig. 3 shows the decay kinetics measured in the Mn-depleted PsbA1/PSII at 432 nm (blue points) and 446 nm (orange points). The data points were averaged from the 5^th^ to the 10^th^ flash when Tyr_D_^●^ was formed in all the centers. With the semi-logarithmic plot used in Panel B of Fig. 3, the decays at both 432 nm and 446 nm were almost linear from 10 ns to 1 µs and had a similar *t*_1/2_ of ~ 200 ns. The P_680_^+^ reduction has been studied earlier in detail in the conditions used (Faller et al. 2001, Rappaport et al. 2009, Sugiura et al. 2010). The *t*_1/2_ value in Fig. 3B is very close to the value of 190 ns found previously (Faller et al. 2001, Rappaport et al. 2009).

The detection of the feature with troughs at 434 nm and 446 nm at times as short as 10-20 ns after the flash shows that it cannot originate from anything other than [P_D1_P_D2_]^+^ or Q_A_^−^ because the reduction of [P_D1_P_D2_]^+^ and the oxidation of Q_A_^−^ are not significant at that time (*t*_1/2_ for [P_D1_P_D2_]^+^ reduction ~200 ns, and for Q_A_^−^ oxidation ~ 400 µs to 1 ms). As mentioned in the introduction, the “W-shape” structure between 440 nm and 460 nm was also observed in spectra after the removal of the Q_A_^−^-*minus*-Q_A_ contribution (Diner et al. 2001). In addition, the decay with a *t*_1/2_ of ~ 200 ns is much too fast to correspond to the forward electron transfer from Q_A_^−^ to Q or Q_A_^−^, nor to charge recombination between Q_A_^−^ and [P_D1_P_D2_]^+^ (~ 1ms). Furthermore, the *t*_1/2_ of ~ 200 ns corresponds well with the kinetics for the electron transfer from Tyr_Z_ or Tyr_D_ to [P_D1_P_D2_]^+^ (Faller et al. 2001). Finally, this negative spectral feature is not observed after the first flash, while Q_A_^−^ is also formed 20 ns after the first flash, confirming that the spectral feature does not arise from Q_A_^−^ itself.

These considerations argue strongly in favor of [P_D1_P_D2_]^+^ rather than Q_A_^−^ being the species responsible for the spectral feature between 440 nm and 460 nm. The question is now whether these absorption changes originate from the [P_D1_P_D2_]^+^ species or if they originate from an electrochromic response of a pigment and/or cofactor induced by the formation of [P_D1_P_D2_]^+^. In order to obtain information on this subject we have recorded the [P_D1_P_D2_]^+^-*minus*-[P_D1_P_D2_] difference spectra in various types of PSII with mutations known to affect some of the cofactors in the vicinity of P_D1_ or P_D2_.

The first of these PSII was the Tyr_D_-less mutant (Sugiura et al. 2004). The experiment in Fig. 2 shows the W-shaped double trough feature at 434 nm and 446 nm is absent when Tyr_D_ is reduced and present when Tyr_D_^●^ is present. It therefore seemed possible that this feature arises from an absorption change in Tyr_D_^●^ generated by the formation of [P_D1_P_D2_]^+^.

Fig. 4 shows the spectra recorded 20 ns after the laser flash illumination in Mn-depleted Tyr_D_-less PSII with PsbA1 as the D1 protein (*i.e.* PsbA1/PsbD1-Y160F PSII) dark adapted for ~ 3-4 h at pH 8.6. The black spectrum was recorded after the first flash and the red spectrum is an average of the ΔI/I measured from the 5^th^ to 10^th^ laser flash illumination. The two spectra are very similar, if not identical, and, surprisingly, they are identical to the [P_D1_P_D2_]^+^-*minus*-[P_D1_P_D2_] difference spectra in the presence of Tyr_D_^●^ and not to those in the presence of Tyr_D_. The blue spectrum that is the red spectrum *minus* the black spectrum is flat around 446 nm.

**Figure 4:**
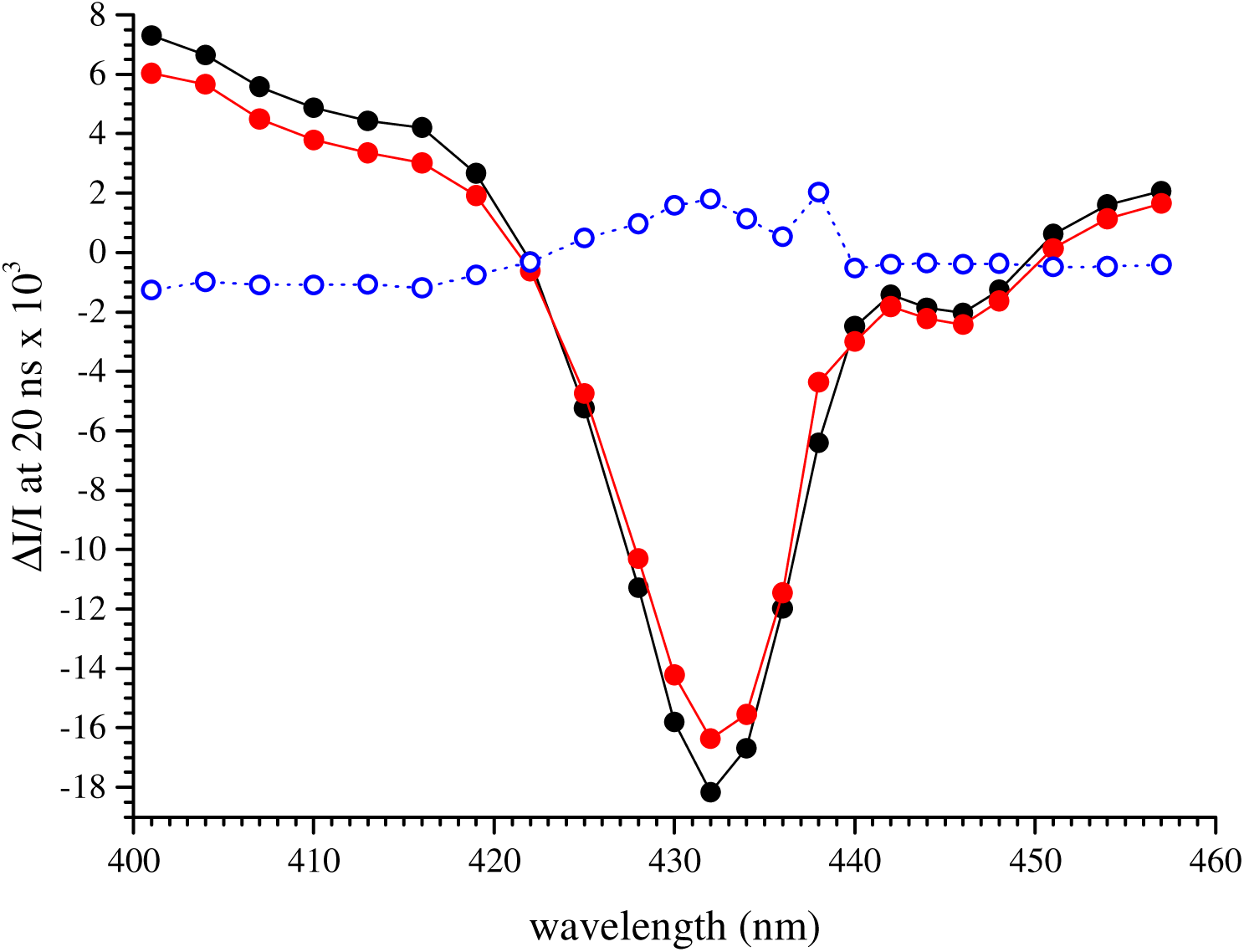
spectra recorded 20 ns after a laser flash illumination in Mn-depleted PsbA1/PsbD1-Y160F PSII dark adapted for 3-4 h at pH 8.6. The black full circles were recorded after the 1^st^ flash and the red full circles are the average the measurements done from the 5^th^ to the 10^th^ 5 ns laser flash. The blue spectrum is the red spectrum *minus* the black spectrum. The Chl concentration was 25 µg mL^−1^ and 100 µM PPBQ was added before the measurements.

It is clear from the results in Fig. 4 that the presence of Tyr_D_^●^ is not required for the detection of the 440 nm to 460 nm double-trough spectral feature and is not directly responsible for this spectral feature. The structure of PSII around Tyr_D_ at pH 8.6 is unknown (see for example Hienerwadel et al. 2008) and the same is true when Tyr_D_ is replaced by phenylalanine in the Tyr_D_-less mutant, however it seems possible that the local electrostatic environment in this mutant could mimic the situation occurring in presence of Tyr_D_^●^. To test the effects of modifications in the H-bond network in this region, the spectra were recorded in two other mutants.

These two mutants were PsbA1-PsbD1/H189L single mutant (Un et al. 2007), in which the H-bonding histidine partner of Tyr_D_ is absent, and the PsbA1-PsbD1/Y160F-H189L double mutant (Sugiura et al. unpublished) in which both Tyr_D_ and its H-boning histidine partner are absent. In the PsbA1-PsbD1/H189L single mutant, the oxidation of Tyr_D_ occurs with a very low efficiency. Consequently, the spectra after the 1^st^ or following flashes are taken in conditions where Tyr_D_^●^ is not present. The results obtained in these two mutants are shown in Fig. 5. Panel A in Fig. 5 shows the spectra recorded in the PsbA1-PsbD1/H189L single mutant. Panel B shows the spectra recorded in the double mutant PsbA1-PsbD1/Y160F-H189L. The black full circles were recorded after the 1^st^ flash and the red full circles are the average of the measurements done from the 5^th^ to the 10^th^ flash.

**Figure 5:**
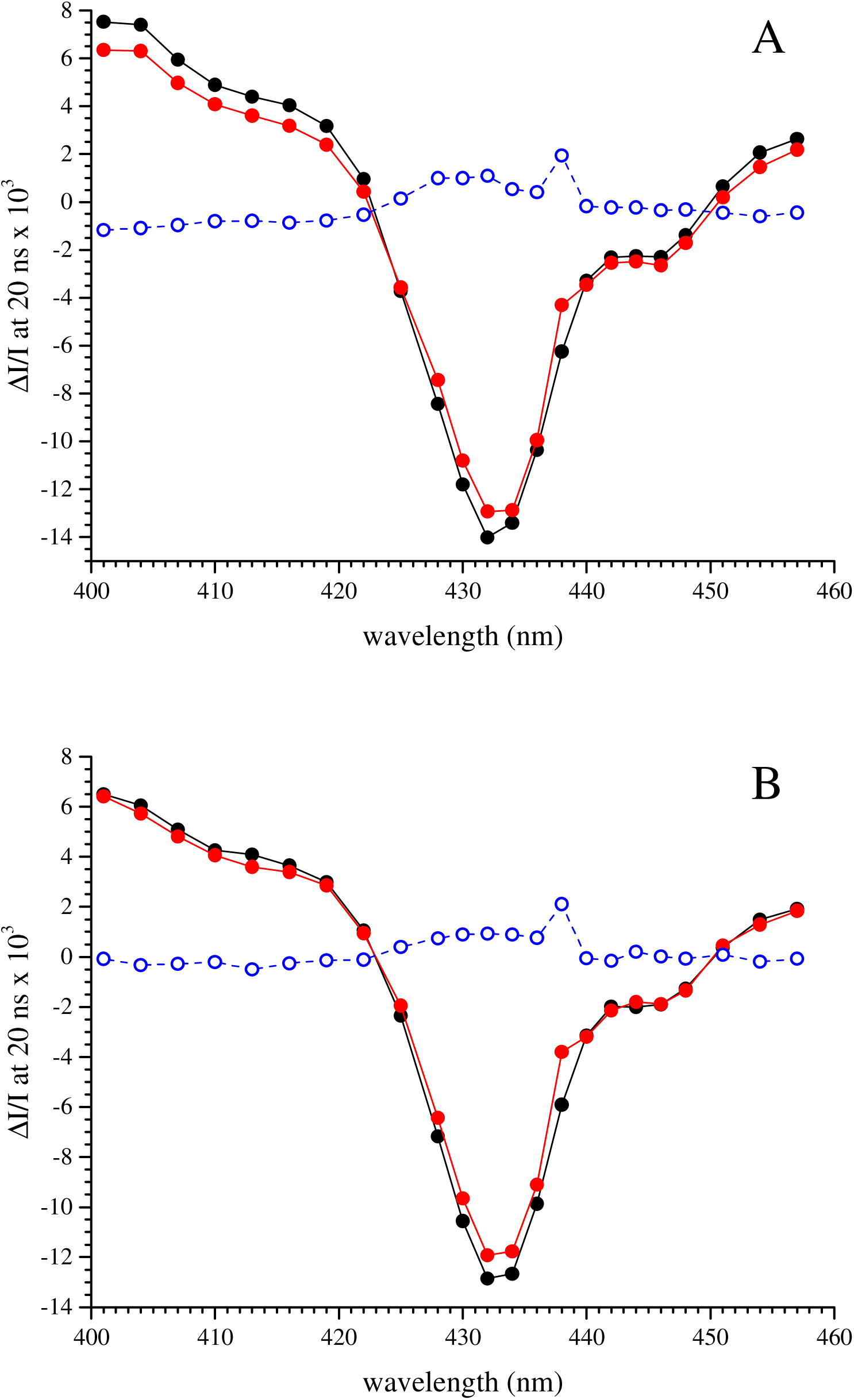
Spectra recorded 20 ns after a laser flash illumination in PsbA1-PsbD1/H189L PSII (Panel A) and in PsbA1-PsbD1/Y160F-H189L (Panel B). The samples were dark adapted for 3-4 h at pH 8.6 before the measurements. The black full circles were recorded after the 1^st^ flash and the red full circles are the average the measurements done from the 5^th^ to the 10^th^ flash. The blue spectra are the red spectra *minus* the black spectra. The Chl concentration was 25 µg mL^−1^ and 100 µM PPBQ was added before the measurements.

In the PsbA1-PsbD1/H189L mutant, the spectral feature that best appears, as a dip at 446 nm, is present when Tyr_D_ is not oxidized whatever the number of flashes given to the sample. This contrasts with the situation in PsbA1-PSII and PsbA3-PSII, where its formation required the presence of Tyr_D_^●^. In the PsbA1-PsbD1/Y160F-H189L double mutant, the modification of the H-bond network due to loss of the Tyr_D_ and its H-bond partner, His 189, had no effect when compared with the PsbA1-PsbD1/Y160F single mutant. It should however be noted that when compared to the situation in PsbA1-PSII and PsbA3-PSII, the spectra between 432 nm and 440 nm appeared slightly broader on the longer wavelength side of the spectrum. In Fig. 5A and 5B, the blue spectra, which are the red spectrum *minus* the black spectrum, are flat around 446 nm.

In the following, we address the situation in other mutants known to modify the spectral properties of P_D1_ and of some of the cofactors around it.

The first of these mutants is the PsbA3/H198Q, in which the His ligand of P_D1_ is replaced by a Gln, see Fig. 1. In this mutant, the [P_D1_P_D2_]^+^-*minus*-[P_D1_P_D2_] difference spectrum is shifted to the blue by ~ 3 nm (Diner et al. 2001, Sugiura et al. 2016). Despite this blue shift in the main bleach, the 440 nm to 460 nm spectral feature remained unaffected both in inactive (*i.e.* Mn-depleted) PSII from *Synechocystis* PCC6803 (Diner et al. 2001) at pH 5.9 and in O_2_ evolving PSII from *T. elongatus* at pH 6.5 (Sugiura et al. 2016), with a trough at 446 nm in these two types of PSII.

The second mutant is the PsbA3/T179H in which the properties of Chl_D1_ are strongly modified (Schlodder et al. 2008, Takegawa et al. 2019). In this mutant, the Qy transition of Chl_D1_ is shifted to the red by ~ 2nm both in inactive PSII from *Synechocystis* PCC6803 (Schlodder et al. 2008) and in O_2_ evolving PSII from *T. elongatus* at pH 6.5 (Takegawa et al. 2019). In the PsbA3/T179H *T. elongatus* mutant, the feature with a trough at 446 nm remained unaffected. However, this negative result can only be taken as inconclusive evidence against a role of Chl_D1_ in forming the spectral feature with the trough at 446 nm.

The third mutant on the D1 side is the PsbA3/E130Q mutant. The residue 130 of PsbA is H-bonded to the 13^1^-keto of Phe_D1_, affecting its midpoint potential and shifting its Qx band in the ~535-545 nm spectral region due to the changing strength of the H-bond (Merry et al. 1998, Sugiura et al. 2014). Fig. 6 shows that the 440 nm to 460 nm spectral feature *i*) remained unaffected in Mn-depleted PsbA3/E130Q PSII at pH 8.6 and *ii*) was absent on the first flash as in Mn-depleted PsbA3 PSII (Fig. 2). The data here show that the mysterious spectral feature in the Soret region does not seem to contain a contribution from a Phe_D1_ bandshift. However, it is not certain that a change in the Qx band of the absorption spectrum of Phe_D1_ would also be accompanied by a change in the Soret region (Sugiura et al. 2014).

**Figure 6:**
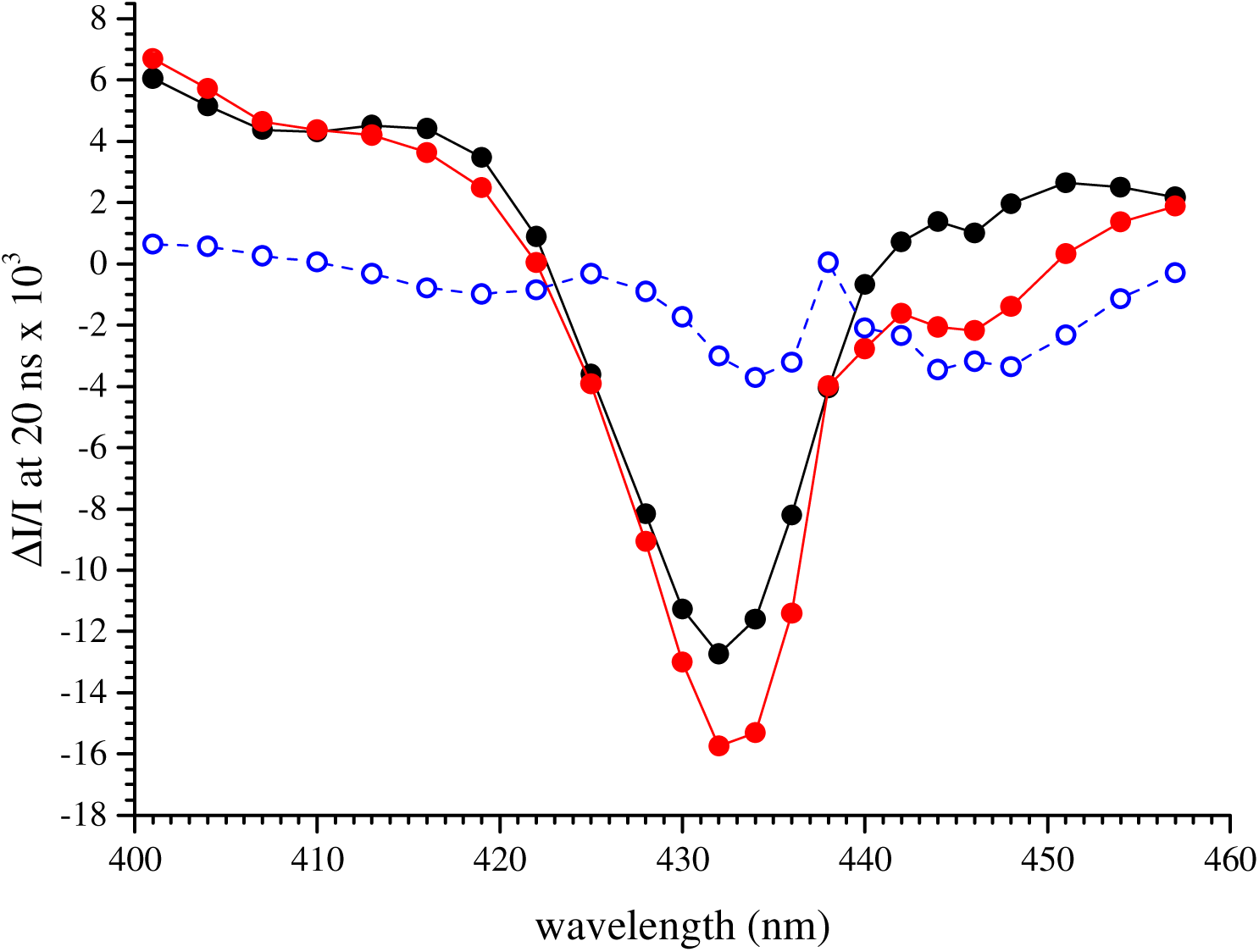
spectra recorded 20 ns after a laser flash illumination in Mn-depleted PsbA3/E130Q PSII, which was dark adapted for 3-4 h at pH 8.6. The black full circles were recorded after the 1^st^ flash and the red full circles are the average the measurements done from the 5^th^ to the 10^th^ flash. The blue spectrum is the red spectrum *minus* the black spectrum. The Chl concentration was 25 µg mL^−1^ and 100 µM PPBQ was added before the measurements.

Probing the modifications on the D2 side is more difficult. By comparing the electrochromic band-shifts of Phe_D1_ and Phe_D2_ in the Q_X_ band region around 550 nm, which are triggered by the formation of either Tyr_Z_^●^ or Tyr_D_^●^ in PsbA1-PSII and PsbA3-PSII, it was shown that the Q_X_ bandshift of Phe_D2_ induced by Tyr_D_^●^ formation was slightly red shifted by 2-3 nm compared to the Phe_D1_ bandshift observed upon the formation of Tyr_Z_^●^. This was interpreted by taking into account, as shown in Fig. 1, that the H-bonded residue to Phe_D1_ is a glutamate (PsbA1/E130) and the H-bonded residue to Phe_D2_ is a glutamine (PsbD/Q129) (Boussac et al. 2020). However, once again the conclusion remains tentative, as a modification of the Qx band does not necessarily imply a change in the Soret region making again this negative result as an inconclusive evidence against a role of Phe_D2_ in the feature at 446 nm.

Fig. 7 shows the spectra recorded in a PsbA3-PsbD1/I178T mutant PSII (Sugiura et al. submitted). The modifications of the Chl_D2_ in this mutant are expected to be comparable to those of the Chl_D1_ in the equivalent D1 mutation, *i.e.*, PsbA3/T179 (Takegawa et al. 2019). Fig. 7 shows again that there is no effect of the PsbD1/I178T mutation on the [P_D1_P_D2_]^+^-*minus*-[P_D1_P_D2_] spectra, and this is taken as an indication that Chl_D2_ is not the origin of the 440 nm to 460 nm double trough absorption feature.

**Figure 7:**
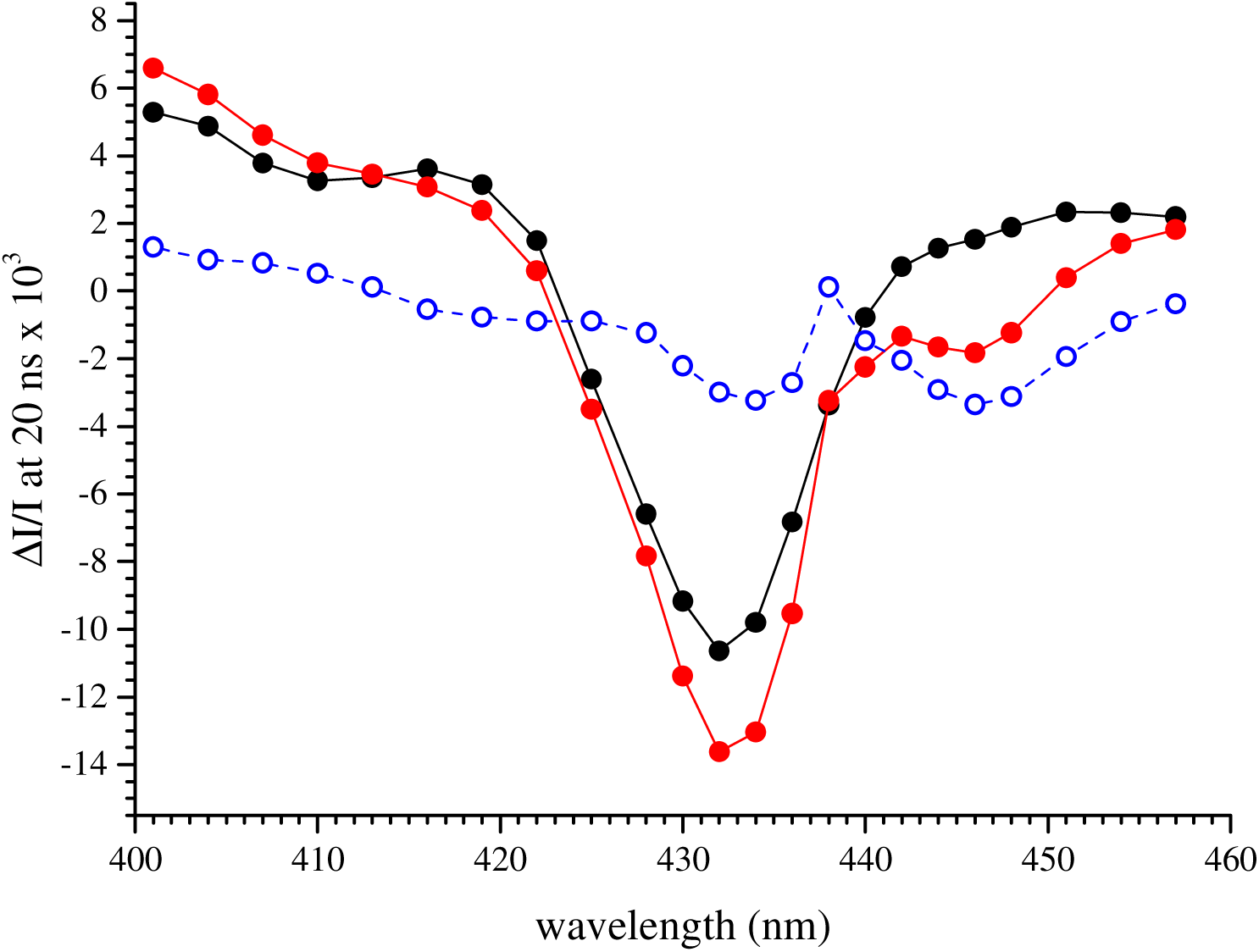
spectra recorded 20 ns after a laser flash illumination in Mn-depleted PsbA3-PsbD1/I178T PSII, which was dark adapted for 3-4 h at pH 8.6. The black full circles were recorded after the 1^st^ flash and the red full circles are the average the measurements done from the 5^th^ to the 10^th^ flash. The blue spectrum is the red spectrum *minus* the black spectrum. The Chl concentration was 25 µg mL^−1^ and 100 µM PPBQ was added before the measurements.

Two other types of modified PSII from *T. elongatus* have been studied. Panel A of Fig. 8, shows that the 43H (His tagged) strain grown in the presence of 3-Fluorotyrosine. In the 43H PSII purified from these culture conditions, the tyrosine residues are replaced by a 3-Fluorotyrosine (Rappaport et al. 2009). Despite that, there was no significant difference in the [P_D1_P_D2_]^+^-*minus*-[P_D1_P_D2_] spectra with respect to the spectra in Fig. 3, recorded with a normal PsbA1-PSII. The second PSII, in Panel B of Fig. 8, is a mutant in which the D1 protein is PsbA3 and in which the heme of Cyt*b*_559_ is lacking (Sugiura et al. 2015). As the heme of Cyt*b*_559_ has a Soret absorption in the spectral region studied here (Kaminskaya et al. 1999), a putative electrochromic band shift of either the reduced or the oxidized form of this heme was a possible candidate as the source of the 440 to 460 nm spectral feature. As it can be seen in Panel B of Fig. 8, this hypothesis can be ruled out since the difference between the red and black spectra is similar to that in PsbA3-PSII.

**Figure 8:**
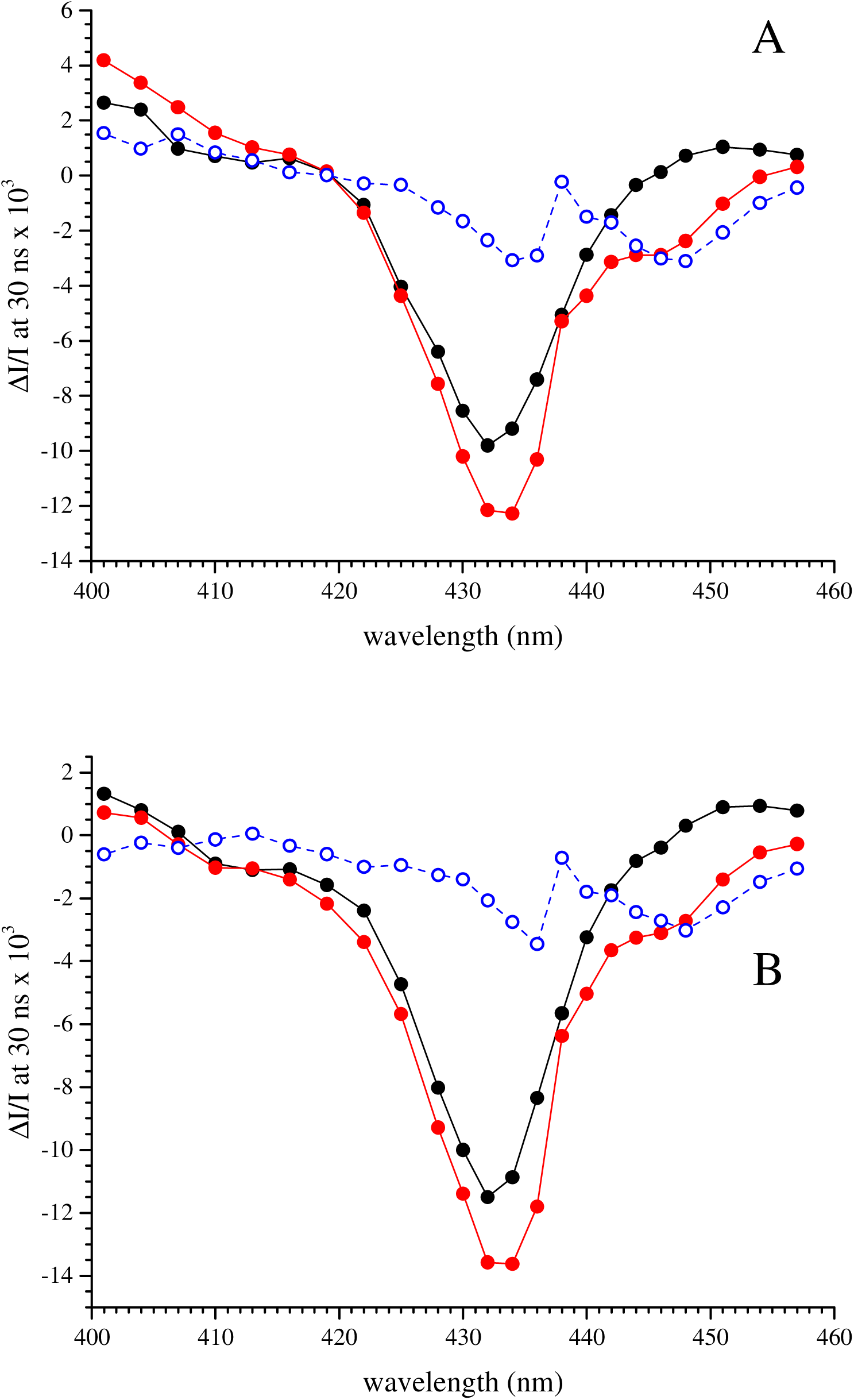
Spectra recorded 30 ns after a laser flash illumination in PsbA1-PSII containing 3-Fluorotyrosine (Panel A) and in the PsbA3/PsbE-H23A PSII (Panel B). The samples were dark adapted for 3-4 h at pH 8.6 before the measurements. The black full circles were recorded after the 1^st^ flash and the red full circles are the average the measurements done from the 5^th^ to the 10^th^ flash. The blue spectra are the red spectra *minus* the black spectra. The Chl concentration was 25 µg mL^−1^ and 100 µM PPBQ was added before the measurements.

The same measurements were made in a PSII variant that has more significant modifications, namely PSII purified from *C. thermalis* grown under far-red light, in which Chl_D1_ is supposed to be a Chl-*d*, or less likely a Chl-*f*, (Nürnberg et al. 2018). Indeed, a recent cryo-EM structure argued for Chl_D1_ being the Chl-*d* in the far-red PSII of *Synechococcus elongatus* PCC7335 (Gisriel et al. 2022). In addition to this Chl-*d/f* in the reaction centre, 4 Chl-*f* (or 1 Chl-*d* and 3 Chl-*f*) replace 4 of the others 34 Chl-*a*. The location of these additional antenna pigments in the far-red PSII of *C. thermalis* has not been determined experimentally yet, but candidates have been proposed based on structural considerations (Nürnberg et al 2018). For this sample, the spectra, shown in Panel A of Fig. 9, were recorded 20 ns after the flashes. The first observation here is that the bleaching induced by the formation of [P_D1_P_D2_]^+^ peaks at the same wavelength (~ 432 nm) as in Chl-*a* only PSII. It should be noted that, in methanol, the Soret band of Chl-*f* is blue-shifted to ~ 400 nm and that of Chl-*d* is red-shifted to ~ 456 nm when compared to Chl-*a*, *e.g.* (Chen 2019). If, as seems likely, some shifts in the absorption occur also when these chlorophyll variants are located in the PSII reaction center, this would indicate that neither Chl-*f* or Chl-*d* would be involved in the P_D1_P_D2_ pair, in agreement with the suggestion of Nurnberg et al. (2018), but discussed by Judd et al. (2020). Nevertheless the fact that the [P_D1_P_D2_]^+^ signal at 432 nm was larger after the 1^st^ than after the followings, the spectral differences between 440 and 460 nm remain clearly unaffected in this PSII with a trough at 446 nm that is observed predominantly after the 5^th^ to 10^th^ flash.

**Figure 9:**
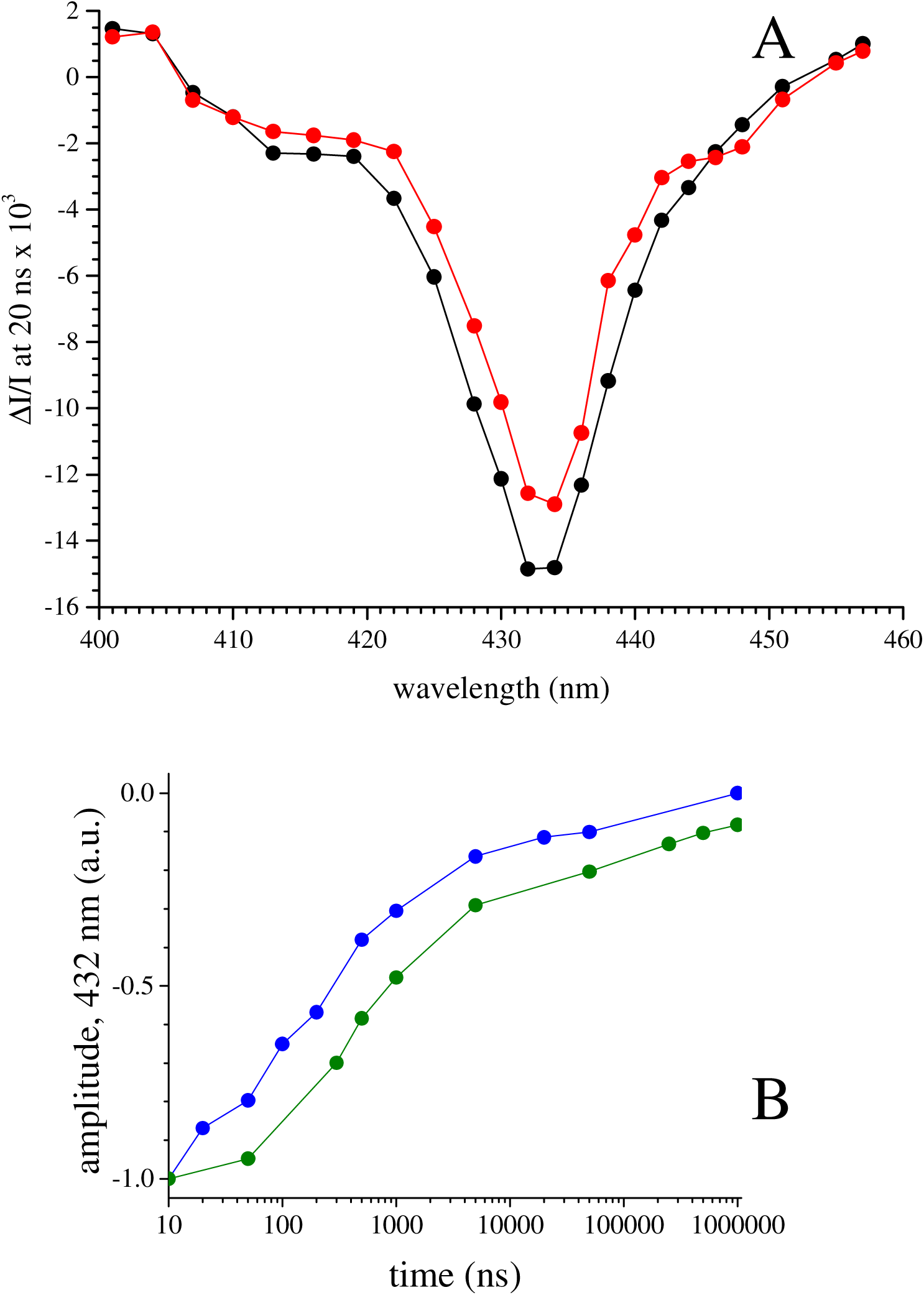
Panel A, spectra recorded 20 ns after each of a series of laser flash illuminations in Mn-depleted PSII purified from *C. thermalis* grown under far red light. The spacing between the 10 flashes of the sequence was 2 s. The sample was dark adapted at room temperature for 3-4 h at pH 8.6 before the recording of the spectra. The black full circles were recorded after the 1^st^ flash and the red full circles are the average the measurements done from the 5^th^ to the 10^th^ flash. The amplitude of the spectra was normalized to a Chl concentration of 25 µg mL^−1^. Panel B, decay kinetics measured at 432 nm from 10 ns to 1 ms in Mn-depleted PsbA1-PSII (blue data points) and in Mn-depleted PSII from *C. thermalis* (green data points). The data points are averaged from the 5^th^ to the 10^th^ flash, *i.e.* when all Tyr_D_^●^ was formed in all centers. The amplitude of the signal at 10 ns was normalized to −1. The Chl concentration was 25 µg mL^−1^ and 100 µM PPBQ was added before the measurements.

Panel B of Fig. 9 compares the decay at 432 nm at pH 8.6 of [P_D1_P_D2_]^+^ in the Mn-depleted PSII from WT*1 *T. elongatus* (blue data points) with that from *C. thermalis* (green data points) In *C. thermalis,* the decay of [P_D1_P_D2_]^+^ occurred with a *t*_1/2_ close to 400 ns (to be compared to the *t*_1/2_ close to 200 ns in PsbA1-PSII from *T. elongatus*). Because the width of the spectrum in Fig. 9A is comparable to that in wild-type PSII from *T. elongatus*, a change in the P_D1_^+^P_D2_ ↔ P_D1_P_D2_^+^ equilibrium seems unlikely to explain the decay rate of [P_D1_P_D2_]^+^ twice as slow in *C. thermalis* than in PsbA1-PSII in *T. elongatus*. It has been discussed that in *C. thermalis* grown under far red light the D1 protein has the highest sequence identity with PsbA3 in *T. elongatus* (Viola et al. 2022). Since in PsbA3-PSII from *T. elongatus* the [P_D1_P_D2_]^+^ decay rate in Mn-depleted PSII and at high pH was slowed down by a factor ~ 2 when compared to PsbA1-PSII (Sugiura et al. 2010) the *t*_1/2_ are similar in *C. thermalis* grown under far red light and PsbA3-PSII in *T. elongatus*. This confirms that the energy levels of the electron transfer cofactors Tyr_Z_ and P_680_ are similar in the two PSII (Viola et al. 2022).

Finally, we have repeated the experiment in the D2/H197A mutant done in *Synechocystis* PCC 6803 (Hayase et al. 2023). In this mutant, the histidine axial ligand of P_D2_ has been replaced by an alanine unable to bind to the Mg^2+^ of P_D2_ (see Fig. 1B) and it has been already studied (Diner et al. 2001, Hayase et al. 2023). Panel A in Fig. 10 shows the spectra recorded 20 ns after the laser flash illumination in Mn-depleted D2/H197A-PSII dark adapted for ~ 3-4 h at pH 8.6. The black spectrum was recorded after the first flash and the red spectrum is an average of the ΔI/I measured from the 5^th^ to 10^th^ laser flash illumination. The two spectra are very similar, if not identical. For comparison, the averaged spectrum from the 5^th^ to the 10^th^ flash and recorded in WT*1-PSII from *T. elongatus* is also shown in blue (open circles). Two main observations can be done.

**Figure 10:**
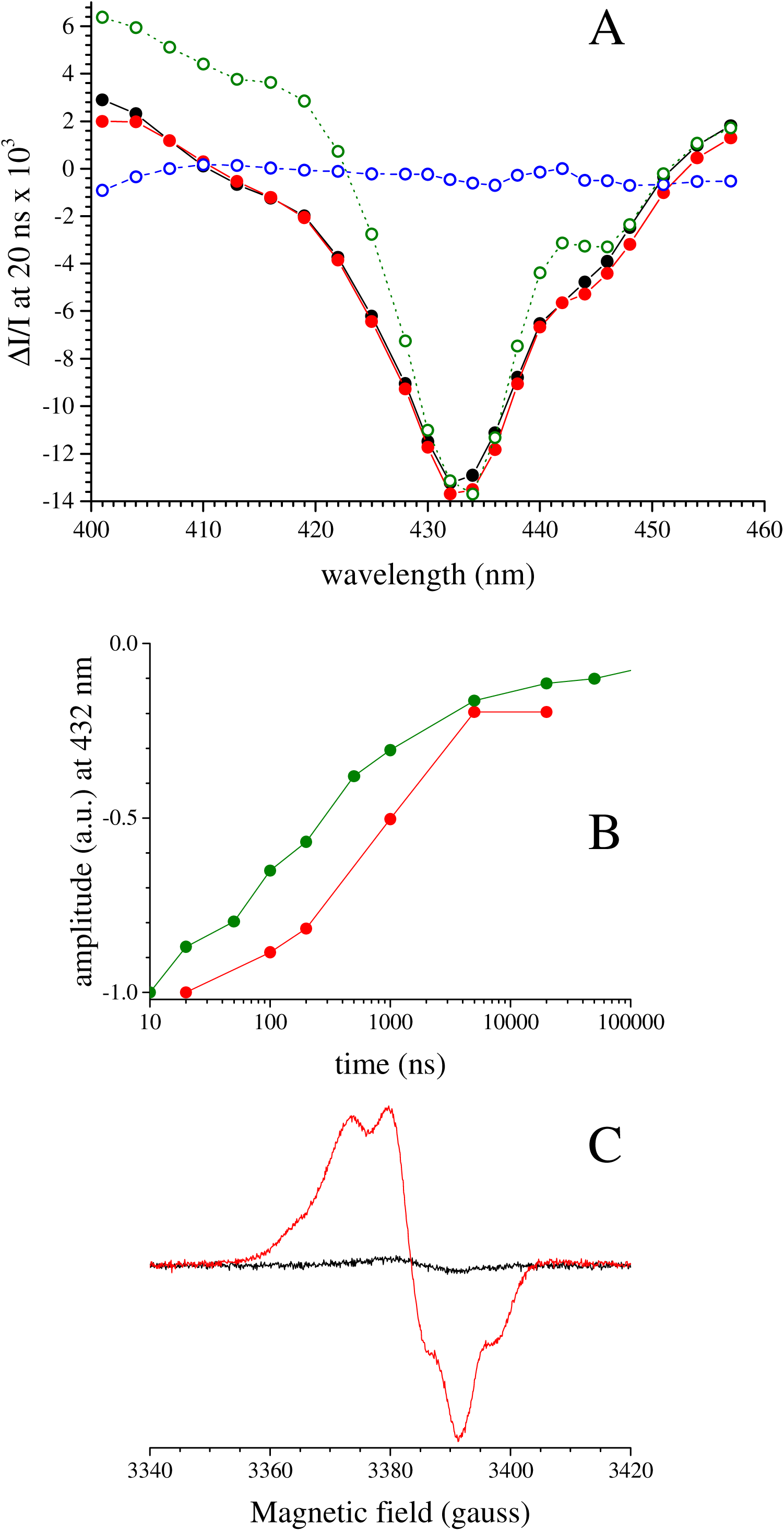
Panel A, spectra recorded 20 ns after a laser flash illumination in Mn-depleted PsbD/H197A PSII from *Synechocystis* 6803, which was dark adapted for 3-4 h at pH 8.6. The black full circles were recorded after the 1^st^ flash and the red full circles are the average the measurements done from the 5^th^ to the 10^th^ flash. The blue spectrum recorded in WT*1-PSII is replotted from Fig. 2. Panel B, decay kinetics at 432 nm in Mn-depleted PsbA1-PSII (orange) and PsbD/H197A (red). The amplitude of the signal at *t* = 0 was normalized to −1. The Chl concentration was 25 µg mL^−1^ and 100 µM PPBQ was added before the measurements in both samples. Panel C, EPR spectra around *g* = 2 in the dark-adapted PsbD/H197A PSII (black spectrum) and after 9 flashes given at room temperature. Instrument settings: temperature 15 K, modulation amplitude, 2.8 G; microwave power, 2 µW; microwave frequency, 9.4 GHz; modulation frequency, 100 kHz.

Firstly, although we cannot say if the [P_D1_P_D2_]^+^Q_A_^−^-*minus*-[P_D1_P_D2_]Q_A_ difference spectrum is red shifted by 1-2 nm as previously mentioned (Diner et al. 2001), the difference spectrum is significantly broader than in WT*1-PSII as previously observed (Diner et al. 2001). Such a broadening could arise from a more delocalized positive charge in [P_D1_P_D2_]^+^. Unfortunately, the broadening of the spectrum does not allow us to detect clearly a shift of the “W-shaped” spectrum. To test the hypothesis of a more delocalized cation, the decay of [P_D1_P_D2_]^+^ has been measured at 432 nm after several flashes, *i.e*. when all Tyr_D_ is expected to be oxidized and the electron donor is only Tyr_Z_. Panel B in Fig. 10 shows the results. The green data points are those recorded in WT*1-PSII and the red data points are those recorded in the D2/H197A-PSII. Clearly, the decay is ~ 5 times slower in the D2/H197A-PSII. This slowdown is not due to a species effect (*Synechocystis vs T. elongatus*) because we have seen that the decay of [P_D1_P_D2_]^+^ (measured at 820 nm) occurred with the same rate (*t*_1/2_ ~ 200 ns) in both species (Faller et al. 2001). The possibility that the cation is more localized on P_D2_ in the D2/H197A is therefore a likely hypothesis. Diner et al. (2001) have estimated that, in this mutant, the difference in the reduction potential between the couples Tyr ^●^/Tyr and [P_D1_P_D2_]^+^/[P_D1_P_D2_] was decreased by 31 mV and that the E*m* of the [P_D1_P_D2_]^+^/[P_D1_P_D2_] couple was decreased by 19 mV. By measuring the thermoluminescence arising for the S Q ^−^/DCMU charge recombination in whole cells we found comparable results (not shown). Such effects of the D2/H197A mutation are expected to slow down the electron transfer from Tyr_Z_ to [P_D1_P_D2_]^+^.

Secondly, and surprisingly, the feature at around 446 nm is already present on the first flash. From what we have observed above either Tyr_D_ would be not oxidizable in this mutant or Tyr_D_^●^ would be present in the dark-adapted PSII. We have therefore recorded the EPR spectra in the dark-adapted D2/H197A-PSII (black spectrum in Panel C of Fig. 10) and after 9 flashes given at room temperatures (red spectrum in Panel C of Fig. 10). Without any ambiguity, Tyr_D_ is not detectable in the dark-adapted PSII and is formed by the flash illumination.

## Conclusion

Wild-type PSII showed a double trough feature in the Soret region of the absorption spectrum when [P_D1_P_D2_]^+^ was formed in the presence of Tyr_D_^●^ (see Fig. 2 blue spectrum). This “W-shaped” signal formed with [P_D1_P_D2_]^+^, with troughs at 434 nm and 446 nm, is not detected on the first flash when Tyr_D_ is reduced and it becomes detectable after the subsequent flashes when Tyr_D_ has been oxidized. From this result, we could expect the “W-shaped” signal to be absent in the Tyr_D_ less mutant, regardless of the number of flashes. Surprisingly, the opposite was observed with the “W-shaped” signal detected after all the flashes. Similarly, the “W-shaped” signal was also formed on all the flashes in the PsbD1/H189L mutant in which Tyr_D_ is present but not oxidable. Finally, the “W-shaped” signal was observed after all the flashes in the PsbD1/H197A mutant in which the residue that coordinates P_D2_ has been changed. These results are consistent with the view that changes in the hydrogen bond network or protein conformation around P_D2_, either in mutants (PsbD1/Y160F, PsbD1/H189L, PsbD/H197A) or upon the oxidation of Tyr_D_, somehow affect either a pigment band, likely a Chl, or the couplings between several pigments. Which pigment and how remain to be elucidated. A range of PSII samples from mutants and variants (as listed in Table 1) were also studied and the double trough feature was seemingly unmodified in all of them. Therefore, there was no evidence for this spectral feature arising from P_D1_, Chl_D1_, Phe_D1_, Phe_D2_, Tyr_Z_, Tyr_D_ and the Cyt*b*_559_ heme. Nevertheless, P_D2_ remains a candidate for the “W-shaped” feature as it is the pigment that is most closely associated with both P_D1_ and Tyr_D_ and the spectroscopic properties of which could be tuned by the H-bond network on the D2-side.

## Abbreviations

Chl: chlorophyll
Chl_D1_/Chl_D2_: monomeric Chl on the D1 or D2 side, respectively
Cyt: cytochrome
DMSO: dimethyl sulfoxide
EPR: Electron Paramagnetic Resonance
P_D1_ and P_D2_: individual Chl on the D1 or D2 side, respectively, which constitute a pair of Chl with partially overlapping aromatic rings (P_680_)
Phe_D1_ and Phe_D2_: pheophytin on the D1 or D2 side, respectively
PPBQ: phenyl *p*–benzoquinone
PSII: Photosystem II
Q_A_: primary quinone acceptor
Q_B_: secondary quinone acceptor
Tyr_D_: the tyrosine 160 of D2 acting as a side-path electron donor of PSII
Tyr_Z_: the tyrosine 161 of D1 acting as the electron donor to P_680_
WT*3: *T. elongatus* mutant strain deleted *psbA_1_* and *psbA_2_* genes and with a His-tag on the carboxy terminus of CP43
WT’: *T. elongatus* mutant strain deleted *psbA_1_, psbA_2_* and *psbD_2_* genes and with a His-tag on the carboxy terminus of CP43
EDTA: Ethylenediaminetetraacetic acid.

## Acknowledgements

This work has been in part supported by (i) the French Infrastructure for Integrated Structural Biology (FRISBI) ANR-10-INBS-05, (ii) the Labex Dynamo (ANR-11-LABX-0011-01), (iii) the JSPS-KAKENHI Grant in Scientific Research on Innovative Areas JP17H064351 and a JSPS-KAKENHI Grant 21H02447 and (iv) the BBSRC grants BB/R001383/1, BB/V002015/1 and BB/R00921X. Adjélé Wilson is thanked for her advice on breaking *Synechocystis* cells using the French press.

## References

Béal D, Rappaport F, Joliot P (1999) A new high-sensitivity 10-ns time-resolution spectrophotometric technique adapted to in vivo analysis of the photosynthetic apparatus. Rev Sci Instrum 70: 202–207. 10.1063/1.1149566

Boussac A, Etienne A-L (1982) Spectral and kinetic pH-dependence of fast and slow Signal-II in Tris-washed chloroplasts. FEBS Lett 148: 113–116. 10.1016/0014-5793(82)81254-4

Boussac A, Sellés J, Sugiura M (2020) What can we still learn from the electrochromic bandshifts in Photosystem II? Biochim Biophys Acta 1861: 148176. 10.1016/j.bbabio.2020.148176

Boussac A, Sugiura M, Rappaport F (2010) Probing the quinone binding site of Photosystem II from *Thermosynechococcus elongatus* containing either PsbA1 or PsbA3 as the D1 protein through the binding characteristics of herbicides. Biochim Biophys Acta 1807: 119–129. 10.1016/j.bbabio.2010.10.004

Brettel K, Schlodder E, Witt HT (1984) Nanosecond reduction kinetics of photooxidized chlorophyll-a (P-680) in single flashes as a probe for the electron pathway, H^+^ release and charge accumulation in the O_2_-evolving complex. Biochim Biophys Acta 766: 403–415. 10.1016/0005-2728(84)90256-1

de Causmaecker S, Douglass JS, Fantuzzi A, Nitschke W, Rutherford AW (2019) Energetics of the exchangeable quinone, Q_B_, in Photosystem II. Proc Natl Acad Sci USA 116: 19458– 19463. 10.1073/pnas.1910675116

Capone M, Sirohiwal A, Aschi M, Pantazis DA, Daidone I (2023) Alternative fast and slow primary charge-separation pathways in Photosystem II. Angew Chem Int Ed 62: e202216276. 10.1002/anie.202216276

Chen M, Schliep M, Willows RD, Cai Z-L, Neilan BA, Scheer H (2010) A Red-shifted chlorophyll. Science 329: 1318–1319. 10.1126/science.1191127

Chen M (2019) Chlorophylls d and f: Synthesis, occurrence, light-harvesting, and pigment organization in chlorophyll-binding protein complexes. Adv Bot Res 90: 121–139. 10.1016/bs.abr.2019.03.006

Conjeaud H, Mathis P (1980) The effect of pH on the reduction kinetics of P-680 in Tris-treatead chloroplasts. Biochim Biophys Acta 590: 353–359. 10.1016/0005-2728(80)90206-6

Cox N, Pantazis DA, Lubitz W (2020) Current understanding of the mechanism of water oxidation in Photosystem II and its relation to XFEL data. Annu Rev Biochem 89: 795–820. 10.1146/annurev-biochem-011520104801

Diner BA, Schlodder E, Nixon PJ, Coleman WJ, Rappaport F, Lavergne J, Vermaas WFJ, Chisholm DA (2001) Site-directed mutations at D1-His198 and D2-His197 of Photosystem II in *Synechocystis* PCC 6803: sites of primary charge separation and cation and triplet stabilization. Biochemistry 24: 9265–9281. 10.1021/bi010121r

Faller P, Debus RJ, Brettel K, Sugiura M, Rutherford AW, Boussac A (2001) Rapid formation of the stable tyrosyl radical in photosystem II. Proc Natl Acad Sci USA 98: 14368–14373. 10.1073/pnas.251382598

Fufezan C, Zhang C-X, Krieger-Liszkay A, Rutherford AW (2005) Secondary quinone in Photosystem II of *Thermosynechococcus elongatus*: Semiquinone-iron EPR signals and temperature dependence of electron transfer. Biochemistry 44: 12780–12789. 10.1021/bi051000k

Gan F, Zhang S, Rockwell NR, Martin SS, Lagarias JC, Bryant DA (2014) Extensive remodeling of a cyanobacterial photosynthetic apparatus in far-red light. Science 345: 1312–1317. 10.1126/science.1256963

Gerken S, Brettel K, Schlodder E, Witt HT (1987) Direct observation of the immediate electron-donor to chlorophyll-a^+^ (P-680^+^) in oxygen-evolving photosystem-complexes – Resolution of manosecond kinetics in the UV. FEBS Lett 223: 376–380. 10.1016/0014-5793(87)80322-8

Giorgi LB, Nixon PJ, Merry SAP, Joseph DM, Durrant JR, Rivas JD, Barber J, Porter G, Klug DR (1996) Comparison of primary charge separation in the photosystem II reaction center complex isolated from wild-type and D1-130 mutants of the cyanobacterium *Synechocystis* PCC 6803. J Biol Chem 271: 2093–2101. 10.1074/jbc.271.4.2093

Gisriel CJ, Wang J, Liu J, Flesher DA, Reiss KM, Huang HL, Yang KR, Armstrong WH, Gunner MR, Batista VS, Debus RJ, Brudvig GW (2022a) High-resolution cryo-electron microscopy structure of Photosystem II from the mesophilic cyanobacterium, Synechocystis sp. PCC 6803. Proc Natl Acad Sci USA 119: e2116765118. 10.1073/pnas.2116765118

Gisriel CJ, Shen GZ, Ho M-Y, Kurashov V, Flesher DA, Wang JM, Armstrong AW, Golbeck JH, Gunner MR, Vinyard DJ, Debus RJ, Brudvig GW, Bryant DA (2022b) Structure of a monomeric photosystem II core complex from a cyanobacterium acclimated to far-red light reveals the functions of chlorophylls d and f. J Biol Chem 298: 101424. 10.1016/j.jbc.2021.101424

Hayase T, Shimada Y, Mitomi T, Nagao R, Noguchi T (2023) Triplet delocalization over the reaction center chlorophylls in Photosystem II. J Phys Chem 127: 1758–1770. 10.1021/acs.jpcb.3c00139

Hienerwadel R, Diner BA, Berthomieu C (2008) Molecular origin of the pH dependence of tyrosine D oxidation kinetics and radical stability in photosystem II. Biochim Biophys Acta 1777: 525–531. 10.1016/j.bbabio.2008.04.004

Holzwarth AR, Müller MG, Reus M, Nowaczyk M, Sander J, Rögner M (2006) Kinetics and mechanism of electron transfer in intact Photosystem II and in the isolated reaction center: pheophytin is the primary electron acceptor. Proc Natl Acad Sci USA 103: 6895–6900. 10.1073/pnas.0505371103

Joliot P, Barbieri G, Chabaud R (1969) A new model of photochemical centers in system 2. Photochem Photobiol 10: 309–329. 10.1111/j.1751-1097.1969.tb05696.x

Judd M, Morton J, Nürnberg D, Fantuzzi A, Rutherford AW, Purchase R, Cox N, Krausz E (2020) The primary donor of far-red photosystem II: Chl(D1) or P-D2? Biochim Biophys Acta 1861: 148248. 10.1016/j.bbabio.2020.148248

Kaminskaya O, Kurreck J, Irrgang KD, Renger G, Shuvalov VA (1999) Redox and spectral properties of cytochrome b(559) in different preparations of photosystem II. Biochemistry 49: 16223–16235. 10.1021/bi991257g

Kok B, Forbush B, McGloin M (1970) Cooperation of charges in photosynthetic O_2_ evolution–I. A linear four step mechanism. Photochem Photobiol 11: 457–475. 10.1111/j.1751-1097.1970.tb06017.x

Lubitz W, Chrysina M, Cox N (2019) Water oxidation in Photosystem II. Photosynth Res 142: 105–125. 10.1007/s11120-019-00648-3

Merry SAP, Nixon PJ, Barter LMC, Schilstra M, Porter G, Barber J, Durrant JR, Klug DR (1998) Modulation of quantum yield of primary radical pair formation in Photosystem II by site-directed mutagenesis affecting radical cations and anions. Biochemistry 37: 17439– 17447. 10.1021/bi980502d

Mirkovic T, Ostroumov EE, Anna JM, van Grondelle R, Govindjee, Scholes GD (2017) Light absorption and energy transfer in the antenna complexes of photosynthetic organisms. Chem Rev 117: 249–293. 10.1021/acs.chemrev.6b00002

Müh F, Zouni A (2005) Extinction coefficients and critical solubilisation concentrations of Photosystems I and II from *Thermosynechococcus elongatus*. Biochim Biophys Acta 1708: 219–228. 10.1016/j.bbabio.2005.03.005

Nakajima Y, Ugai-Amo N, Tone N, Nakagawa A, Iwai M, Ikeuchi M, Sugiura M, Suga M, Shen J-R (2022) Crystal structures of photosystem II from a cyanobacterium expressing *psbA2* in comparison to *psbA3* reveal differences in the D1 subunit. J Biol Chem 298: 102668. 10.1016/j.jbc.2022.102668

Nurnberg DJ, Morton J, Santabarbara S, Telfer A, Joliot P, Antonaru LA, Ruban AV, Cardona T, Krausz E, Boussac A, Fantuzzi A, Rutherford AW (2018) Photochemistry beyond the red limit in chlorophyll f-containing photosystems. Science 360: 1210–1213. 10.1126/science.aar8313

Ogami S, Boussac A, Sugiura M (2012) Deactivation processes in PsbA1-Photosystem II and PsbA3-Photosystem II under photoinhibitory conditions in the cyanobacterium *Thermosynechococcus elongatus*. Biochim Biophys Acta 1817: 1322–1330. 10.1016/j.bbabio.2012.01.015

Rappaport F, Boussac A, Force DA, Peloquin J, Brynda M, Sugiura M, Un S, Britt RD, Diner BA (2009) Probing the coupling between proton and electron transfer in Photosystem II core complexes containing a 3-fluorotyrosine. J Am Chem Soc 131: 4425–4433. 10.1021/ja808604h

Renger G (2012) Mechanism of light induced water splitting in Photosystem II of oxygen evolving photosynthetic organisms. Biochim Biophys Acta 1817: 1164–1176. 10.1016/j.bbabio.2012.02.005

Romero E, Novoderezhkin VI, van Grondelle R (2017) Quantum design of photosynthesis for bio-inspired solar-energy conversion. Nature 543: 355–365. 10.1038/nature22012

Rutherford AW, Boussac A, Faller P (2004) The stable tyrosyl radical in Photosystem II: why D? Biochim Biophys Acta 1655: 222–230. 10.1016/j.bbabio.2003.10.016

Schlodder E, Renger T, Raszewski G, Coleman WJ, Nixon PJ, Cohen RO, Diner BA (2008) Site-directed mutations at D1-Thr179 of photosystem II in *Synechocystis* sp PCC 6803 modify the spectroscopic properties of the accessory chlorophyll in the D1-branch of the reaction center. Biochemistry 47: 3143–3154. 10.1021/bi702059f

Sedoud A, Cox N, Sugiura M, Lubitz W, Boussac A, Rutherford AW (2011) The semiquinone-iron complex of Photosystem II: EPR signals assigned to the low field edge of the ground state doublet of Q_A_^●^-Fe^2+^ and Q_B_^●^-Fe^2+^. Biochemistry 50: 6012–6021. 10.1021/bi200313p

Shevela D, Kern JF, Govindjee G, Messinger J (2023) Solar energy conversion by photosystem II: principles and structures. Photosynth Res 156: 279–307. 10.1007/s11120-022-00991-y

Suga M, Akita F, Hirata K, Ueno G, Murakami H, Nakajima Y, Shimizu T, Yamashita K, Yamamoto M, Ago H, Shen J-R (2015) Native structure of Photosystem II at 1.95 angstrom resolution viewed by femtosecond X-ray pulses. Nature 517: 99–103. 10.1038/nature13991

Sugiura M, Azami C, Koyama K, Rutherford AW, Rappaport F, Boussac A (2014) Modification of the pheophytin redox potential in *Thermosynechococcus elongatus* Photosystem II with PsbA3 as D1. Biochim Biophys Acta 1837: 139–148. 10.1016/j.bbabio.2013.09.009

Sugiura M, Boussac A, Noguchi T, Rappaport F (2008) Influence of Histidine-198 of the D1 subunit on the properties of the primary electron donor, P680, of Photosystem II in *Thermosynechococcus elongatus*. Biochim Biophys Acta 1777: 331–342. 10.1016/j.bbabio.2008.01.007

Sugiura M, Inoue Y (1999) Highly purified thermo-stable oxygen-evolving photosystem II core complex from the thermophilic cyanobacterium *Synechococcus elongatus* having his-tagged CP43. Plant Cell Physiol 40: 1219–1231. 10.1093/oxfordjournals.pcp.a029510

Sugiura M, Kato Y, Takahashi R, Suzuki H, Watanabe T, Noguchi T, Rappaport F, Boussac A (2010) Energetics in Photosystem II from *Thermosynechococcus elongatus* with a D1 protein encoded by either the *psbA1* or *psbA3* gene. Biochim Biophys Acta 1797: 1491–1499. 10.1016/j.bbabio.2010.03.022

Sugiura M, Nakamura M, Koyama K, Boussac A (2015) Assembly of oxygen-evolving Photosystem II efficiently occurs with the apo-Cytb559 but the holo-Cytb559 accelerates the recovery of a functional enzyme upon photoinhibition. Biochim Biophys Acta 1847: 276–285. 10.1016/j.bbabio.2014.11.009

Sugiura M, Osaki Y, Rappaport F, Boussac A (2016) Corrigendum to “Influence of Histidine-198 of the D1 subunit on the properties of the primary electron donor, P680, of Photosystem II in *Thermosynechococcus elongatus*”. Biochim Biophys Acta 1857: 1943–1948. 10.1016/j.bbabio.2016.09.012

Sugiura M, Rappaport F, Brettel K, Noguchi T, Rutherford AW, Boussac A (2004) Site-directed mutagenesis of the *Thermosynechococcus elongatus* photosystem II: The O_2_-evolving enzyme lacking the redox-active tyrosine D. Biochemistry 43: 13549–13563. 10.1021/bi048732h

Takegawa Y, Nakamura M, Nakamura S, Noguchi T, Sellés J, Rutherford AW, Boussac A, Sugiura M (2019) New insights on Chl_D1_ function in Photosystem II from site-directed mutants of D1/T179 in *Thermosynechococcus elongatus*. Biochim Biophys Acta 1860: 297–309. 10.1016/j.bbabio.2019.01.008

Un S, Boussac A, Sugiura M (2007) Characterization of the Tyrosine-Z radical and Its Environment in the Spin-Coupled S_2_Tyr ^●^ State of Photosystem II from *Thermosynechococcus elongatus*. Biochemistry 46: 3138–3150. 10.1021/bi062084f

Viola S, Roseby W, Santabarbara S, Nürnberg D, Assunção R, Dau H, Sellés J, Boussac A, Fantuzzi A, Rutherford AW (2022) Impact of energy limitations on function and resilience in long-wavelength Photosystem II. eLife 11: e79890. 10.7554/eLife.79890

